# Off-target detection of CRISPR-Cas9 nuclease *in vitro* with CROFT-Seq

**DOI:** 10.1101/2025.02.17.638614

**Authors:** Paulius Toliusis, Algirdas Grybauskas, Tomas Sinkunas, Tautvydas Karvelis, Giedrius Sasnauskas, Mindaugas Zaremba

## Abstract

Programmable CRISPR-Cas9 nucleases have become invaluable tools for genome editing. However, off-target cleavage by these nucleases could lead to unintended changes in the edited genome. Detection of off-target sites is critical to make genome editing technology safe and predictable. Although current *in vitro* methods for off-target detection can identify these sites, they are time-consuming, complex, and relatively costly. Here, we present CROFT-Seq (**CR**ISPR nuclease **of**f-**t**arget detection by **seq**uencing), a sensitive, rapid, and cost-effective assay for the genome-wide detection of Cas9 off-target sites *in vitro*. CROFT-Seq performs comparably to the common currently used *in vitro* methods and serves as a valuable and efficient tool for the rapid assessment of genome-editing nuclease specificity. Notably, a high proportion of the top-ranked off-targets identified by CROFT-Seq were validated in cells, highlighting its effectiveness as a predictor of off-target sites.

## Introduction

Programmable genome editing nucleases, such as CRISPR-Cas9, hold great promise for therapeutic applications and have attracted widespread attention from the scientific community(1–3). These enzymes generate double-strand breaks (DSBs) in genomic DNA that can be repaired in the cell by error-prone non-homologous end joining (NHEJ) that can result in insertion/deletion mutations (indels) or precise homology-directed repair (HDR)(4,5). Nevertheless, unintentional cleavage can occur at various DNA sites similar to the target sequence, where even rare instances of cleavage may result in unwanted or potentially deleterious gene mutations and translocations(6).

The most widely used programmable nuclease for clinical use is Cas9 from *Streptococcus pyogenes*(7). The Cas9 protein associates with guide RNA (gRNA) composed of either crRNA-tracrRNA or single-guide RNA and forms an active ribonucleoprotein (RNP) complex(8,9). This complex can be directed to any genomic site of interest based on complementarity between gRNA and target DNA. Recognition of the target DNA also depends on a short protospacer adjacent motif (PAM) located downstream of the target site, which is recognized by the Cas9 protein. This leads to complementary hybridization of the gRNA to the 20-base pair (bp) target sequence(10). Upon target binding, the HNH and RuvC nuclease domains of the Cas9 protein cleave both DNA strands, generating a double-strand break (DSB). However, the Cas9 complex can bind and cleave similar DNA target sites (off-targets) with nucleotide mismatches, affecting the precision of the programmable genome editing nucleases(11).

A number of genome-wide cell-based and cell-free methods have been developed to detect off-target cleavage sites. The most widely used cell culture-based assays, such as GUIDE-Seq(12), IDVL(13), LAM-HTGTS(14), BLESS/BLISS(15,16), DISCOVER-Seq(17), and ChIP-Seq(18), can directly or indirectly detect Cas9-induced DSBs in the cell making them valuable tools for preliminary evaluations before initiating costly *in vivo* genome editing experiments. However, these methods have several limitations, including high false negative rates, low sensitivity, dependence on cell transfection efficiency, chromatin accessibility, cell type, and other factors(19). Although the cell-free methods detect off-targets only *in vitro*, they detect a significantly higher number of off-target sites and allow precise control of RNP concentration, amount of purified DNA, and cleavage time. This capability enables efficient and rapid RNP pre-screening to identify the most suitable RNP candidates for genome engineering applications. To date, several cell-free genome-wide off-target identification assays have been developed, including Digenome-seq(20), SITE-Seq(21), CIRCLE-Seq/CHANGE-Seq(22,23), CLEAVE-Seq(24), and RGEN-Seq(25). While these techniques can identify a significantly greater number of potential off-target sites than cell-based approaches, they also have certain limitations. For example, Digenome-seq requires a large number (>400 million) of reads per sample due to sequencing of a high background of randomly sheared DNA(20). SITE-Seq, CIRCLE-Seq, RGEN-Seq, and CLEAVE-Seq are relatively sensitive but are prone to false-positive off-target detection (low off-target validation rate)(21,22,24,25). Furthermore, these methods are labor-intensive, time-consuming, and costly and typically require DNA shearing and multiple DNA purification steps using spin columns or magnetic beads, leading to significant DNA loss.

In this study, we present CROFT-Seq, an *in vitro* method designed to thoroughly detect genome-wide off-target sites induced by programmable genome editing nucleases, such as CRISPR-Cas9. We show here that CROFT-Seq exhibits sensitivity comparable to the widely used *in vitro* methods CIRCLE-Seq and SITE-Seq. Furthermore, CROFT-Seq outperforms existing *in vitro* off-target detection methods in terms of cost and hands-on time and is a streamlined, one-tube method offering an easily automated workflow for the entire protocol. Finally, we have shown that CROFT-Seq detects off-target sites at frequencies that closely match those observed in cells, making it an attractive method for benchmarking Cas9 proteins prior to their use for genome editing applications.

## Materials and methods

### Cell culture and transfection

HEK293T cells (ATCC CRL-3216) were cultured in DMEM medium supplemented with 10% FBS (Sigma), 100 U/ml penicillin, and 100 μg/ml streptomycin (Thermo Fisher Scientific) at 37°C and 5% CO_2_. The cells were maintained at confluency below 80%. Cells were plated in a 12-well plate with approximately 250,000 cells per well in 500 μL culture medium one day before transfection. All transfections were performed with 3.6 μL TurboFect transfection reagent (Thermo Fisher Scientific) per 1.4 μg of plasmid encoding only Cas9 or Cas9 and gRNA in 180 μL of DMEM medium. The transfected cells were collected 3 days post-transfection. Genomic DNA was purified with GeneJET Genomic DNA Purification Kit (Thermo Fisher Scientific) and quantified by Qubit 4.0 fluorimeter (Thermo Fisher Scientific).

### CROFT-Seq library preparation

CROFT-Seq experiments with *FANCF, VEGFA1*, and *XRCC5* targeting gRNAs (**Supplementary Protocol**) or without gRNAs were performed on human genomic DNA (Roche). Genomic DNA was treated with Thermosensitive Alkaline Phosphatase (Thermo Fisher Scientific). *In vitro* cleavage reactions were performed in a 40 µl reaction volume containing 100 nM SpCas9, 200 nM gRNA (Synthego), and 1 µg of human genomic DNA. Digested products were ligated to a biotinylated adapter and treated with Exonuclease I (Thermo Fisher Scientific). Ligated DNA products were immobilized on streptavidin-coated MyOne C1 magnetic beads (Thermo Fisher Scientific). The complementary DNA strand was removed with NaOH. The non-biotinylated DNA strand was synthesized using DNA oligonucleotide and T4 DNA polymerase (Thermo Fisher Scientific). DNA was then removed from the magnetic beads and amplified by PCR using Phusion Plus DNA polymerase (Thermo Fisher Scientific). The prepared libraries were quantified on Agilent 2100 Bioanalyzer (Agilent) and sequenced with 100 bp paired-end reads on an Illumina NextSeq 550 or 2000 platform. The depth of coverage was ∼1-10 million reads per sample.

### rhAmpSeq deep sequencing

HEK293T cells were transfected with plasmids expressing Cas9 and gRNAs, as described above. Off-target lists for each gRNA were identified by CROFT-Seq, and the top 5% of off-targets, calculated from the rank 1 off-target site of the read depth, were selected for validation. A total of 90 off-target sites (including on-target sites) were selected for two target sites (*FANCF* and *VEGFA1*) (**Supplementary Tables S3** and **S4**). PCR rhAmpSeq primers were successfully designed for all 90 sites and ordered from Integrated DNA Technologies (IDT) (**Supplementary Table S2**). Following the manufacturer’s protocol, rhAmpSeq PCR libraries were amplified from 25 ng of purified HEK293T genomic DNA duplicates using the rhAmpSeq CRISPR library kit (IDT). The resulting PCR products were pooled into different libraries corresponding to the different gRNAs used for transfection and purified using Ampure XP magnetic beads (Agencourt). The resulting rhAmpSeq libraries were quantified and analyzed on Agilent 2100 Bioanalyzer (Agilent) and sequenced with 100 bp paired-end reads on a NextSeq 2000 instrument. The depth of coverage was approximately 30 million reads per sample (approx. 300,000 reads per off-target).

The sequencing data were analyzed using the rhAmpSeq CRISPR analysis tool (https://eu.idtdna.com/pages/tools/rhampseq-crispr-analysis-tool). For each off-target site, indel frequencies (Δ indels, %) were calculated by subtracting two average control cells (edited with SpCas9 only) from two average transfected cells (edited with SpCas9-gRNA complex). The off-target site was counted as validated if the calculated Δ indel frequency was > 0.1%.

### CROFT-Seq data analysis pipeline

The analysis started with paired-end reads that were used as initial data. The removal of residual adapter sequences and low-quality reads was performed using AdapterRemoval(26) utilizing default parameters and appropriate adapter pairs specified in the method. The cleaned reads were aligned to the genome using BWA-MEM(27) and sorted and indexed with SAMtools(28). The alignment file generation and manipulation were executed using default settings. For each cleavage site search experiment, there were three target and three control samples. Initially, each of the six samples was analyzed independently using the ‘find-cleavage-patterns.jl’ script. This script utilizes a 4 bp reading window to search for read start positions in the aligned R1 reads. If the number of read starts within the reading window range is greater than or equal to 20, the region is included for further analysis. From each included region, the cleavage position is selected based on the presence of reads with forward and reverse directions, or if absent, the position with the highest number of read starts. Subsequently, the target sequence is searched around the chosen cleavage position using the Needleman–Wunsch algorithm(29) (realized in the package ‘BioJulia/BioAlignments.jl’), and the best potential alignment position for the sequence is recorded in the BED format file. After all six samples were analyzed individually, the results were combined using the ‘combine-cleavage-patterns.jl’ script. The target and control samples were combined separately. This combining process aimed to determine if the identified cleavage sites were consistently found within their respective sample groups. The cleavage site is considered to be found if the exact cleavage position is present in all three target samples but not in control samples. Following the combinations, the cleavage sites found in the control samples were excluded from the target samples. The deduction process was performed using the same script as in the combination step. The final list of potential on/off-target regions is output in the BED format file.

## Results

### Development and optimization of CROFT-Seq

Here we aimed to develop an *in vitro* off-target detection method that enhances multiple aspects of the off-target detection process while maintaining high sensitivity. We hypothesized that a one-pot method, without requiring physical or enzymatic DNA fragmentation, would be the most suitable assay for rapid and cost-effective genome-wide detection of off-target sites induced by CRISPR-Cas9 nucleases *in vitro* (**Figure 1, Supplementary Figure S1**, and **Supplementary Protocol**). To achieve these objectives, we implemented and extensively optimized various reaction parameters. Before nuclease treatment, CROFT-Seq utilizes dephosphorylation (treatment with phosphatase) to prevent randomly abundant DNA ends from adapter ligation, thereby increasing sensitivity(24). Next, phosphatase-treated DNA is incubated with CRISPR-Cas9 nuclease, which produces cleavage products predominantly with blunt-end 5’ phosphate containing termini(8,9). To verify cleavage activity, on-target cleavage is assessed using qPCR, by comparing the amount of uncleaved DNA in the reaction treated with the SpCas9-gRNA complex to that in the negative-control reaction, which is treated with SpCas9 alone (**Supplementary Figure S1**). The cleavage products are then ligated to a biotinylated adapter. To minimise DNA loss during the unligated adapter removal step, we designed a unique adapter that can be selectively removed by DNA exonuclease in the same reaction tube. The adapter-ligated DNA is then immobilised on streptavidin-coated magnetic beads to facilitate buffer exchange for subsequent reactions. Next, to reduce the amount of non-specific DNA in the downstream steps, the complementary, non-biotinylated DNA strand is removed by NaOH treatment(30). The remaining immobilized ssDNA is then used as a template to synthesize new complementary, non-biotinylated DNA strands. To avoid the need for DNA fragmentation, CROFT-seq utilizes T4 DNA polymerase along with a primer containing a 12-nucleotide random sequence at the 3’-end, generating short DNA fragments suitable for Illumina sequencing. The DNA is then released from streptavidin-coated magnetic beads, and quantified by qPCR. Finally, DNA is amplified by PCR using oligonucleotides containing Illumina sequencing adapters, purified, and sequenced (**Supplementary Figure S1**).

**Figure 1.**
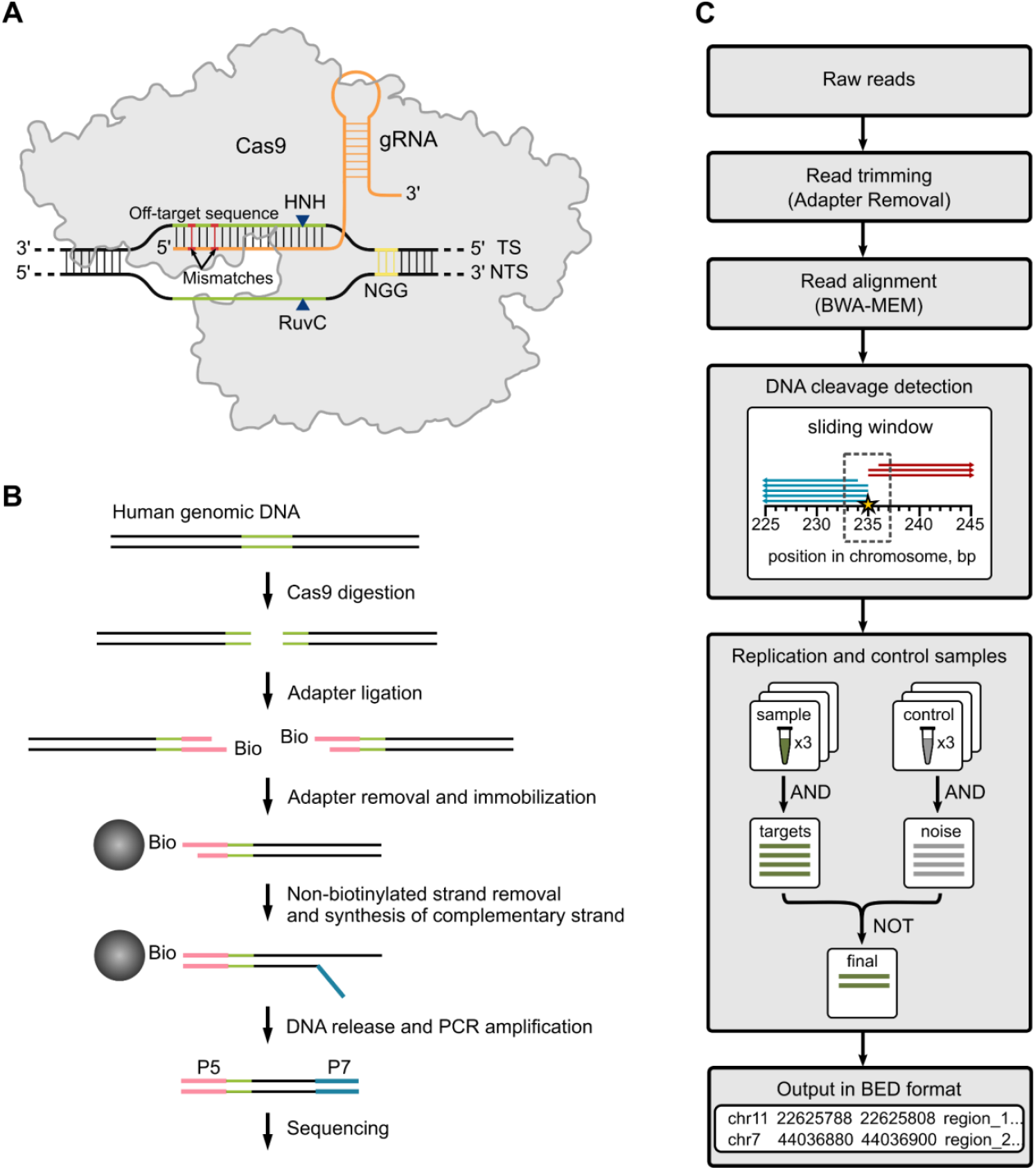
A schematic overview of CROFT-Seq. (**A**) A schematic illustration of *S. pyogenes* Cas9 with a gDNA (orange) bound to an off-target dsDNA (green) containing mismatches (red) proximal to an NGG PAM sequence (yellow). (**B**) Simplified schematic representation of the CROFT-Seq workflow. Human genomic DNA is treated with a phosphatase before digestion with the Cas9 nuclease. The resultant DNA ends are then selectively ligated to a biotinylated adapter. Excess of the adapter is then removed, and the ligated DNA is enriched with magnetic beads. The complementary, non-biotinylated DNA strand is removed, and a new second DNA strand is synthesized. The resultant DNA is released from the beads and amplified by PCR for sequencing. (**C**) Workflow of the CROFT-Seq bioinformatic analysis. Pair-end reads, sequenced and cleaned of residual adapter sequences, are first aligned to the reference genome. Aligned reads are then analyzed with an off-target detection script that searches for steep read depth changes using a 4 bp reading window and prioritizes bidirectionality of the potential off-target-related reads and target sequence similarity. Only the off-target positions detected in all three sample replicates are analyzed further. These positions are filtered out if they are found in all three control samples.

To identify the cleavage sites, we developed an algorithm to analyze the sequencing results. Briefly, sequencing reads are trimmed to remove adapter sequences and then aligned to the reference genome. The aligned reads are then analyzed using the CROFT-Seq pipeline, which searches for read start positions using a 4 bp reading window approach. Potential off-target site positions are identified if they appear in all replicates and are filtered out if they are detected in control reactions (**Figure 1C**).

### Off-target identification by CROFT-Seq

The CROFT-Seq method was tested on human genomic DNA treated with SpCas9 RNP complexes, targeting FANCF, VEGFA1, and XRCC5 sites that have been previously used by other *in vitro* off-target detection methods. A total of 427, 385, and 263 off-target sites were identified for FANCF, VEGFA1, and XRCC5, respectively. Off-targets were ranked by the count of the mapped reads. As expected, most detected off-target sites showed sequence similarity to the on-target site (**Supplementary Table S1**). Furthermore, we generated sequence logos for off-target sites for the FANCF and VEGFA1 guide RNAs, taking into account nucleotide frequency at each position (**Figure 2A**). For both guides, nucleotide frequencies correlated well within the PAM and seed sequences(31,32). This finding aligns with prior studies, including SITE-Seq(21), indicating that mismatches at the PAM and seed regions are less tolerated than mismatches at the PAM-distal end of off-targets.

**Figure 2.**
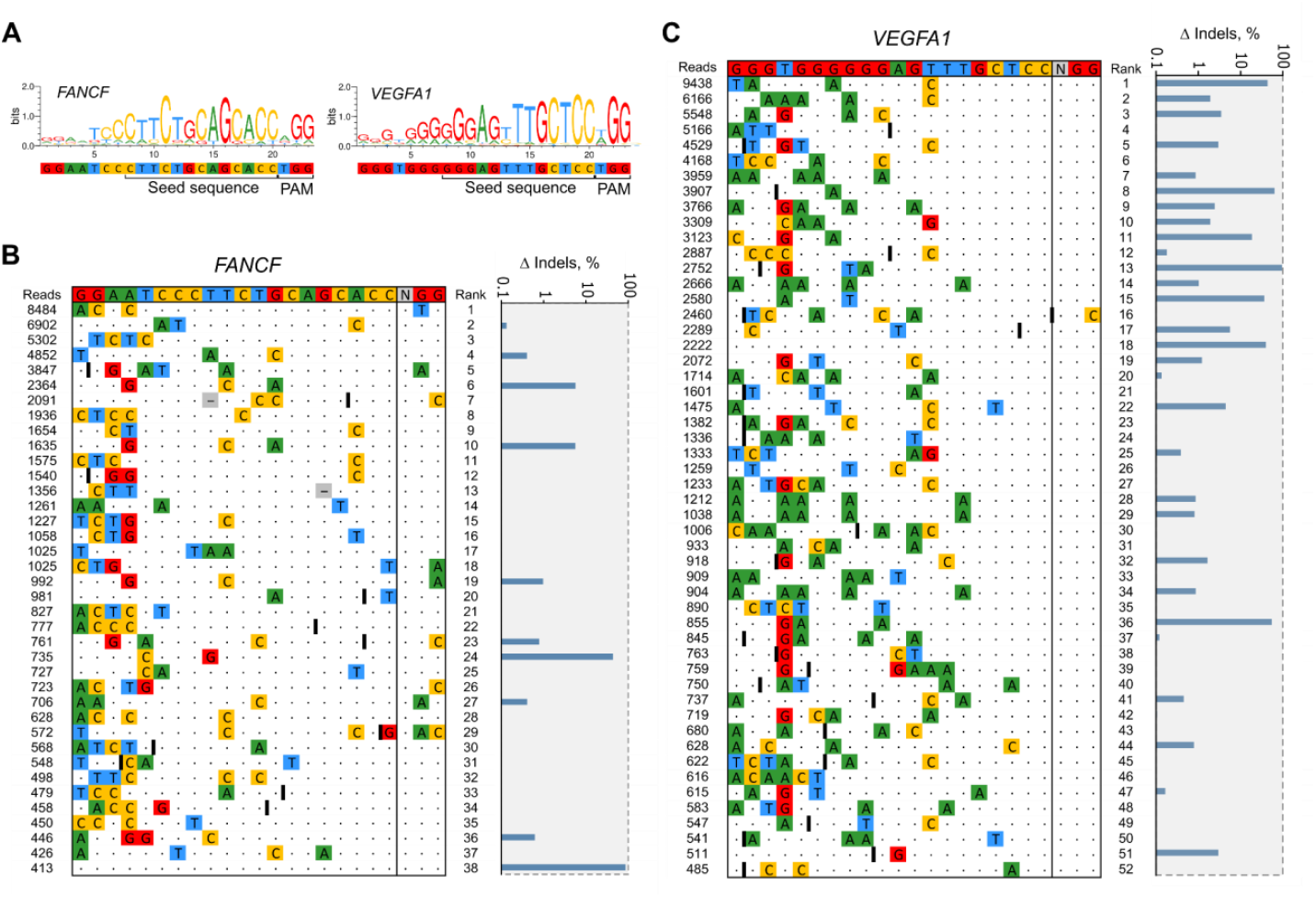
Validation of off-target sites detected by CROFT-Seq. (**A**) Sequence logos generated from FANCF and VEGFA1 off-target sequences detected by CROFT-Seq. Validation results of off-target sites detected by CROFT-Seq for gDNA targeting *FANCF* (**B**) and *VEGFA1* (**C**). The off-target sites whose read counts exceed 5% of the top1 off-target were selected for validation. The on-target site is shown at the top of the off-target list; mismatched nucleotides are colored, gRNA bulges are shown as gray squares with horizontal lines, and DNA bulges are shown as black lines (**I**). On the right is the logarithmic plot showing the changes in indel frequency observed by rhAmpSeq between nuclease-treated and control samples. Off-target sites are considered valid if the change in indel frequency is greater than 0.1%.

We evaluated the reproducibility of CROFT-Seq by comparing read counts between two technical replicates using gRNAs targeting the *FANCF, VEGFA1*, and *XRCC5* sites. The off-target site read counts for these gRNAs were highly correlated, with Pearson R^2^ values of 0.9963, 0.9805, and 0.952, respectively (**Supplementary Figure S2A-C, Supplementary Table S1**). Additionally, we assessed the read count reproducibility between two independent CROFT-Seq reactions (each averaged from three technical replicates) targeting the *FANCF* site, which also demonstrated high reproducibility (R^2^ = 0.9904) (**Supplementary Figure S2D, Supplementary Table S1**).

### *In vivo* validation of off-target sites identified by CROFT-Seq

We further tested whether the off-target sites identified by CROFT-Seq *in vitro* were also cleaved by SpCas9 in cells. HEK293T cells were transfected with plasmids encoding SpCas9 and gRNA targeting the *FANCF* and *VEGFA1* sites. After 3 days post-transfection, the frequencies of insertion/deletion mutations (indels) at the respective sites were quantified using rhAmpSeq (**Supplementary Table S2**). For validation experiments, the off-target sites with read counts exceeding 5% of the number of reads for the most abundant off-target (top1) were selected. This included 38 off-target sites for *FANCF* and 52 for *VEGFA1* (including on-target sites) detected by CROFT-Seq, which were subsequently selected for validation (**Figure 2B, C** and **Supplementary Table S3**). Sites were annotated as edited if the difference in indel frequency (Δ indels, %) between samples treated with SpCas9 and gRNA (cleaved) and those treated with SpCas9 only (control) exceeded a threshold of 0.1%. Validation experiments demonstrated that 10 of the 38 off-target sites for *FANCF* and 30 of the 52 off-target sites for *VEGFA1* were successfully confirmed as edited in HEK293T cells (**Figure 2B, C** and **Supplementary Table S4**).

### Comparison of CROFT-Seq with other *in vitro* off-target detection methods

To measure the performance of CROFT-Seq, we compared data for three different gRNAs, targeting *VEGFA1, FANCF*, and *XRCC5* sites in the human genome that had previously been tested by two of the most widely used *in vitro* off-target detection methods, SITE-Seq and CIRCLE-Seq(21,22) (**Figure 3A, Supplementary Tables S1** and **S5**).

**Figure 3.**
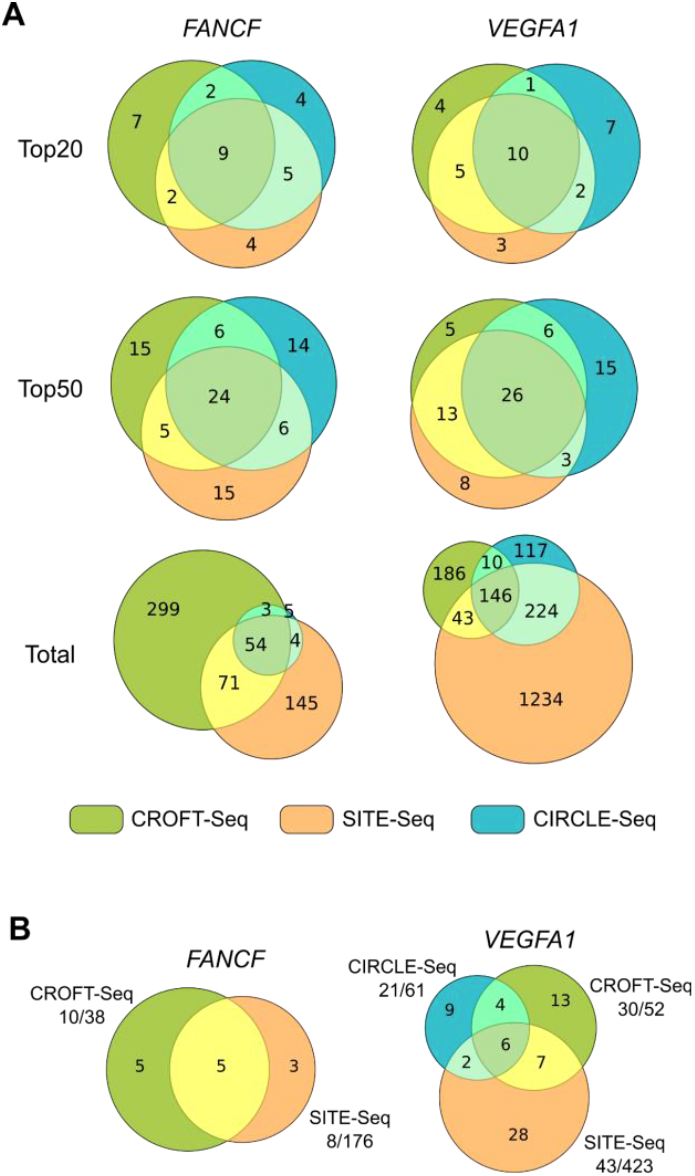
Comparison of CROFT-Seq with SITE-Seq and CIRCLE-Seq for off-target detection. (**A**) Venn diagrams showing the overlap of detected off-target sites between CROFT-Seq, CIRCLE-Seq, and SITE-Seq. The used gRNA targeting *FANCF* or *VEGFA1* is shown at the top, and overlapping ranges (top20, top50 and the whole off-target list) used for comparison are shown on the left. (**B**) Venn diagrams showing overlap of in cells-validated off-target sites for *FANCF* and *VEGFA1* target sites between CROFT-Seq, SITE-Seq and CIRCLE-Seq. The numbers shown after each off-target method name (e. g., 10/38) indicate the total number of off-target sites subjected to validation (38), of which ten (10) off-target sites were validated (Δ indels% > 0.1 - for CROFT-Seq and SITE-Seq, tag integration% > 0 - for CIRCLE-Seq).

For the *FANCF* and *VEGFA1* sites, the top20 and top50 off-target lists of all three CROFT-Seq, SITE-Seq, and CIRCLE-Seq methods overlapped by approx. 50%. The CROFT-Seq and SITE-Seq top50 lists overlapped by approx. 40% of the same *XRCC5* target site (**Supplementary Figure S3A, Supplementary Tables S1** and **S5**). These results indicate that all these methods perform with comparable accuracy in off-target detection.

In addition, we performed a more comprehensive comparison of CROFT-Seq with four *in vitro* off-target detection methods, CIRCLE-Seq, SITE-Seq, RGEN-Seq, and DIGENOME-Seq on *FANCF* and *VEGFA1* sites (**Supplementary Figure S3B, C, Supplementary Tables S1** and **S5**). In this case, the overlap of off-target sites is visualized in an UpSet plot showing the intersections of the methods in the matrix table. CROFT-Seq showed one of the best intersections with other methods, consistent with accurate and robust off-target site detection. Next, we sought to compare how the *in vivo* validated off-target sites of CROFT-Seq overlap with the off-target sites validated by SITE-Seq and CIRCLE-Seq methods (**Figure 3B**). The results show that the validated off-targets of CROFT-Seq overlap well with those of SITE-Seq and CIRCLE-Seq. Moreover, when compared to these methods, the CROFT-Seq showed a high yield of validated off-target sites, suggesting that CROFT-Seq is a sensitive and accurate predictor for off-target sites in cells.

### Comparison of CROFT-Seq with the *in vivo* GUIDE-Seq method

We also compared the overlap of *in vitro* detected and validated off-target sites between CROFT-Seq and the widely used *in vivo* off-target identification method, GUIDE-Seq (**Figure 4A, B**).

**Figure 4.**
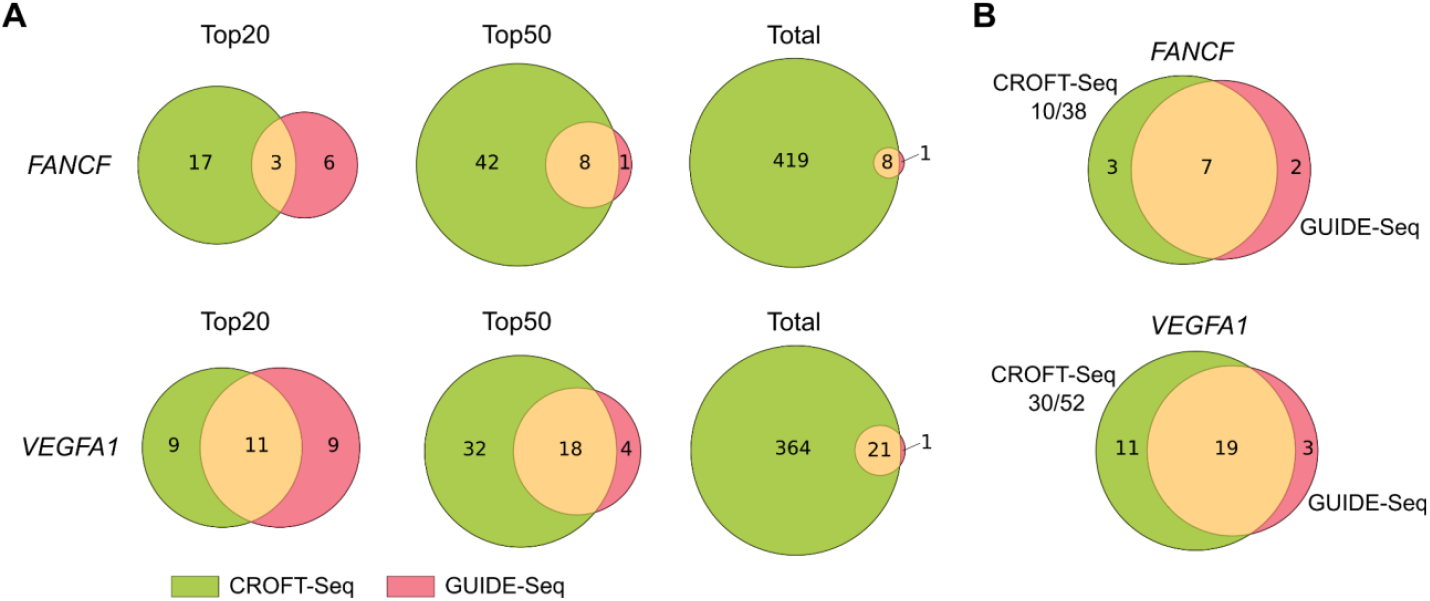
Comparison of CROFT-Seq with the *in vivo* off-target method GUIDE-Seq. (**A**) Venn diagrams showing the overlap at different ranges (top20, top50 and total off-target list) of detected off-target sites between CROFT-Seq and GUIDE-Seq using gRNA targeting the *FANCF* and *VEGFA1* sites. (**B**) Venn diagrams showing the overlap of validated off-target sites for the *FANCF* and *VEGFA1* target sites between CROFT-Seq and GUIDE-Seq. The numbers shown for CROFT-Seq (e.g., 10/38) indicate the total number of off-target sites subjected to validation (38), of which ten (10) off-target sites were validated (Δ indels% > 0.1).

CROFT-Seq detects almost all off-target sites identified by GUIDE-Seq, and most of them are already detected at the top of the CROFT-Seq off-target list (**Figure 4A**, see top50). Careful inspection of the off-targets detected by GUIDE-Seq but undetected by CROFT-Seq revealed that one FANCF off-target was not detected by CROFT-Seq because the number of reads (15) at the cleavage site in one reaction replicate was below the set off-target detection threshold (≥20 reads in each reaction replicate) (**Supplementary Figure S4A**). Another undetected *VEGFA1* off-target was excluded from the CROFT-Seq list by bioinformatic pipeline due to not unambiguously mapped reads at the cleavage site (**Supplementary Figure S4B**). The validated CROFT-Seq off-targets overlapped quite well with the GUIDE-Seq off-targets (**Figure 4B**). Moreover, CROFT-Seq determined additional *in vivo* validated off-targets that were not detected by the GUIDE-Seq (**Figure 4B**). These results demonstrate the power of CROFT-Seq to accurately detect off-target sites occurring *in vivo*.

## Discussion

Genome editing tools such as CRISPR-Cas9 are widely used in clinical trials and in the development of biotechnologically relevant products. While they can be precisely directed to specific genomic DNA sites, they are not always accurate and may lead to unintended genome edits(33–35). Therefore, before selecting and using these tools *in vivo*, it is crucial to verify their efficiency and accuracy to ensure they cleave on-target with minimal off-target edits. *In vitro* methods for off-target detection are particularly well-suited for this purpose. Here, we present CROFT-Seq, our *in vitro* off-target detection method for the comprehensive evaluation of Cas9 cleavage sites in human genomic DNA. CROFT-Seq has several key improvements over other *in vitro* methods (**Table 1 and Supplementary Table S6**).

**Table 1.** Comparison of CROFT-Seq with other *in vitro* off-target detection methods.

| Method | Genomic DNA input, μg | Hands-on time, min | Total time, h | Price per sample, Eur | Reads per sample, mln | Suitable for robotization |
| --- | --- | --- | --- | --- | --- | --- |
| CROFT-Seq | 1 | 45 | 8 | 7 | ~2-5 | Fully (easy) |
| CLEAVE-Seq | 3 | 73 | 12 | 28 | ~5 | Partially (moderate) |
| SITE-Seq | 10 | 80 | 38 | 51 | ~0.62-2.46 | Fully (difficult) |
| CIRCLE-Seq | 25 | 154 | 36 | 66 | ~4-5 | Fully (very difficult) |
| CHANGE-Seq | 1 | 130 | 30 | 70 | ~4-5 | Fully (very difficult) |
| Digenome-Seq | 8 | 261 | 16 | 41 | ~400 | Partially (easy) |
| RGEN-Seq | 3 | 52 | 8 | 75 | ~2 | Fully (difficult) |

CROFT-Seq requires a relatively low amount of genomic DNA (1 μg), making it an effective tool for detecting nuclease-induced off-target sites in cells and tissues where the amount of isolated DNA is limited. In addition, we have significantly reduced both the cost per sample and the hands-on time required to perform CROFT-Seq compared to other methods. This is achieved using individual enzymes, reagents, and buffers with known compositions rather than relatively expensive commercial kits with proprietary and undisclosed compositions used in other methods. Moreover, due to its unique reaction compositions and conditions, CROFT-Seq does not require multiple DNA library purification steps or DNA fragmentation, making it suitable for performing the whole protocol in a single reaction tube, thus allowing for simple automation.

Despite the simple and inexpensive design, CROFT-Seq performs comparably to SITE-Seq and CIRCLE-Seq, the most commonly used *in vitro* off-target detection methods. In our comparison with SITE-Seq and CIRCLE-Seq, we observed that all methods detected the most common *in vitro*-generated off-target sites (**Figure 3**). Furthermore, the CROFT-Seq data analysis algorithm does not rely on additional filtering steps for off-target site identification, such as setting specific MAPQ scores for read alignment or applying mismatch count thresholds for off-target calling. Such filters may introduce bias and negatively impact the overall off-target detection. Lastly, our *in vivo* validation results indicate that CROFT-Seq is a reliable predictor of *in cellulo* off-target sites due to its high success rate in off-target validation (**Figure 2**).

Specific optimization of the CROFT-Seq assay for off-target site detection means that the on-target sites may not always rank at the top of the off-target list. Indeed, the FANCF, VEGFA1, and XRCC5 on-target sites were ranked at positions 40, 18, and 35 of the off-target lists, respectively. The ability to detect on-target sites is highly dependent on the concentration of RNP complex, which can influence their ranking(21). Moreover, the RuvC nuclease domain of SpCas9 can trim the non-target strand after cleavage(36), making the cleavage products less effectively ligatable to the blunt-ended adapter. Additionally, certain off-target sites might be cleaved more rapidly than on-target sites due to sequence context or the secondary structure of the target DNA(37,38). Consequently, these off-target sites are likely to be detected with higher read counts compared to on-target sites.

Next, we compared CROFT-Seq with the cell-based method GUIDE-Seq, which is a widely used approach for direct detection of off-target sites in the edited cells. The results showed that the off-target sites detected by CROFT-Seq match well with those identified by the GUIDE-Seq method. Furthermore, CROFT-Seq has detected additional validated off-target sites that were not identified by GUIDE-Seq (**Figure 4**). However, a major drawback of cell-free methods is their tendency to produce a significant number of false-positive off-target sites, which are challenging to eliminate(39). Addressing this issue, we still require extensive algorithm optimization or the development of alternative *in vitro* off-target detection methods.

The low cost, time efficiency, and ease of automation for high throughput studies make CROFT-Seq a powerful and attractive tool for identifying off-target sites of genome editing nucleases. Finally, we envisage that CROFT-Seq could be adapted for the off-target detection of other emerging genome editing tools, such as Cas12, TnpB, and base editors.

## Data availability

CROFT-Seq and rhAmpSeq sequencing raw data can be accessed at NCBI BioProject database under accession number PRJNA1156337. Other supporting data of this study is available from the corresponding author upon request. CROFT-Seq open-source software is free and available at https://github.com/agrybauskas/croft-seq-analysis.

## Supplementary data

Supplementary Data are available at NAR Online.

## Acknowledgments

We thank prof. V. Siksnys for discussions and suggestions. Also, we thank R. Zedaveinyte for her support with *in vivo* transfection experiments. We thank the EMBL GeneCore facility for performing high-throughput sequencing. *Author contributions:* M.Z., G.S., T.K., and T.S. designed experiments to develop the CROFT-Seq method. P.T. performed all CROFT-Seq and cell-based validation experiments. A.G. wrote the CROFT-Seq bioinformatic analysis pipeline. P.T., A.G., T.K., G.S., and M.Z. contributed to developing the CROFT-Seq bioinformatic algorithm and sequencing data analysis. P.T. and M.Z. wrote the manuscript with the help of other co-authors.

