## Supplementary Figures for "Off-target detection of CRISPR-Cas9 nuclease *in vitro* with CROFT-Seq"

### Supplementary data

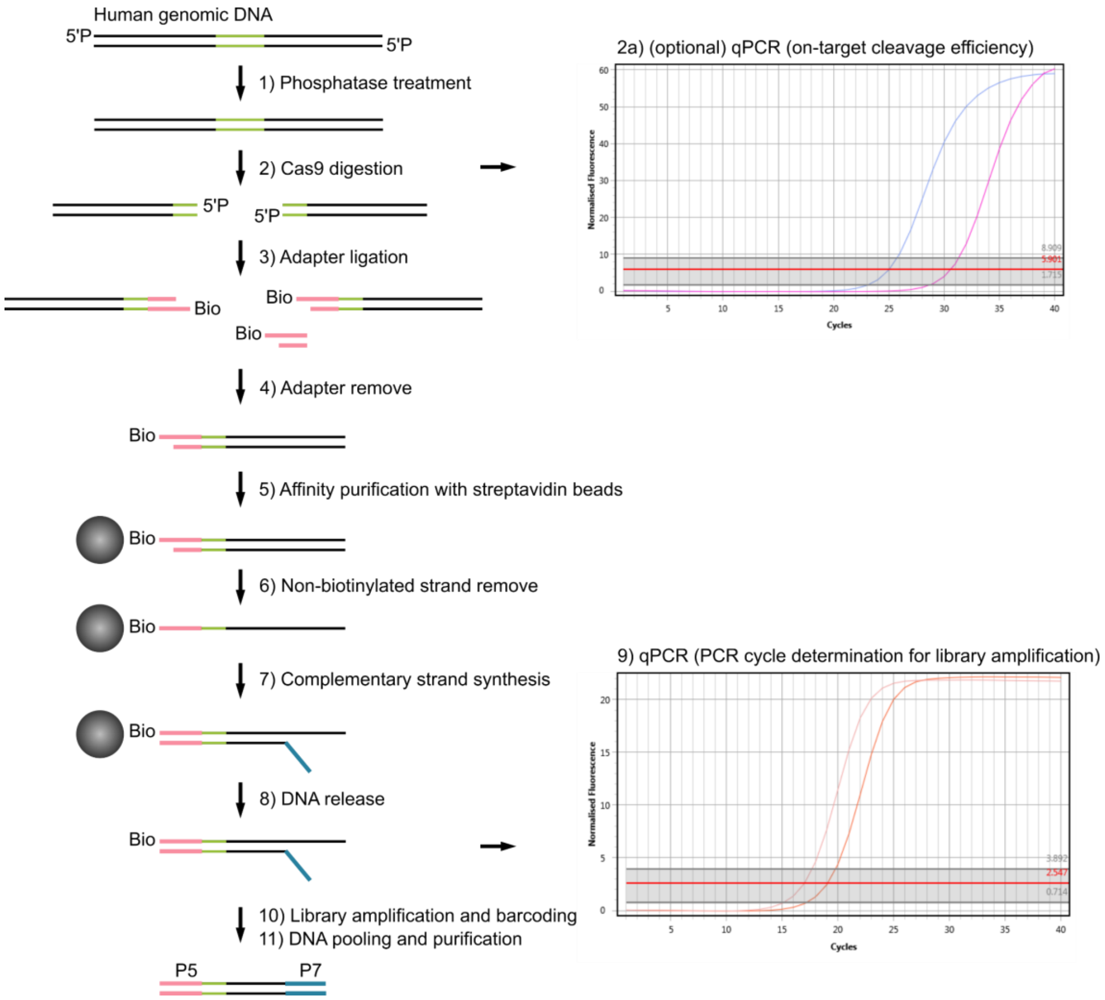


**Supplementary Figure S1.** Detailed schematic overview of CROFT-Seq. Human genomic DNA is treated with Thermosensitive Alkaline Phosphatase, which reduces free DNA ends containing phosphates. The DNA is then treated with a genome-editing nuclease, such as Cas9. In this step, the on-target cleavage efficiency is also measured by qPCR. Representative qPCR amplification curves of DNA samples treated with Cas9 only (blue) or Cas9 with *FANCF* gRNA (purple) are shown on the top right. After DNA cleavage, the products are selectively ligated to the biotinylated adapter. The residual adapter is then removed with DNA exonuclease I. The ligated DNA products were then immobilized on streptavidin-coated MyOne C1 magnetic beads. The non-biotinylated DNA strand is removed with NaOH under conditions required for DNA strand separation. The immobilized ssDNA is then used for complementary strand synthesis using oligonucleotide (containing a 12 N nucleotide sequence where N is A, G, C, or T) and T4 DNA polymerase. The resulting DNA is then released from streptavidin-coated magnetic beads, and the number of PCR cycles required for library amplification is determined by qPCR. Representative qPCR amplification curves of DNA samples treated with Cas9 only (orange) or Cas9 with *FANCF* gRNA (light orange) are shown on the bottom right. The DNA library is then amplified with Phusion Plus DNA polymerase using DNA oligonucleotides with Illumina barcode sequences. Individual DNA libraries are pooled and purified using magnetic beads.

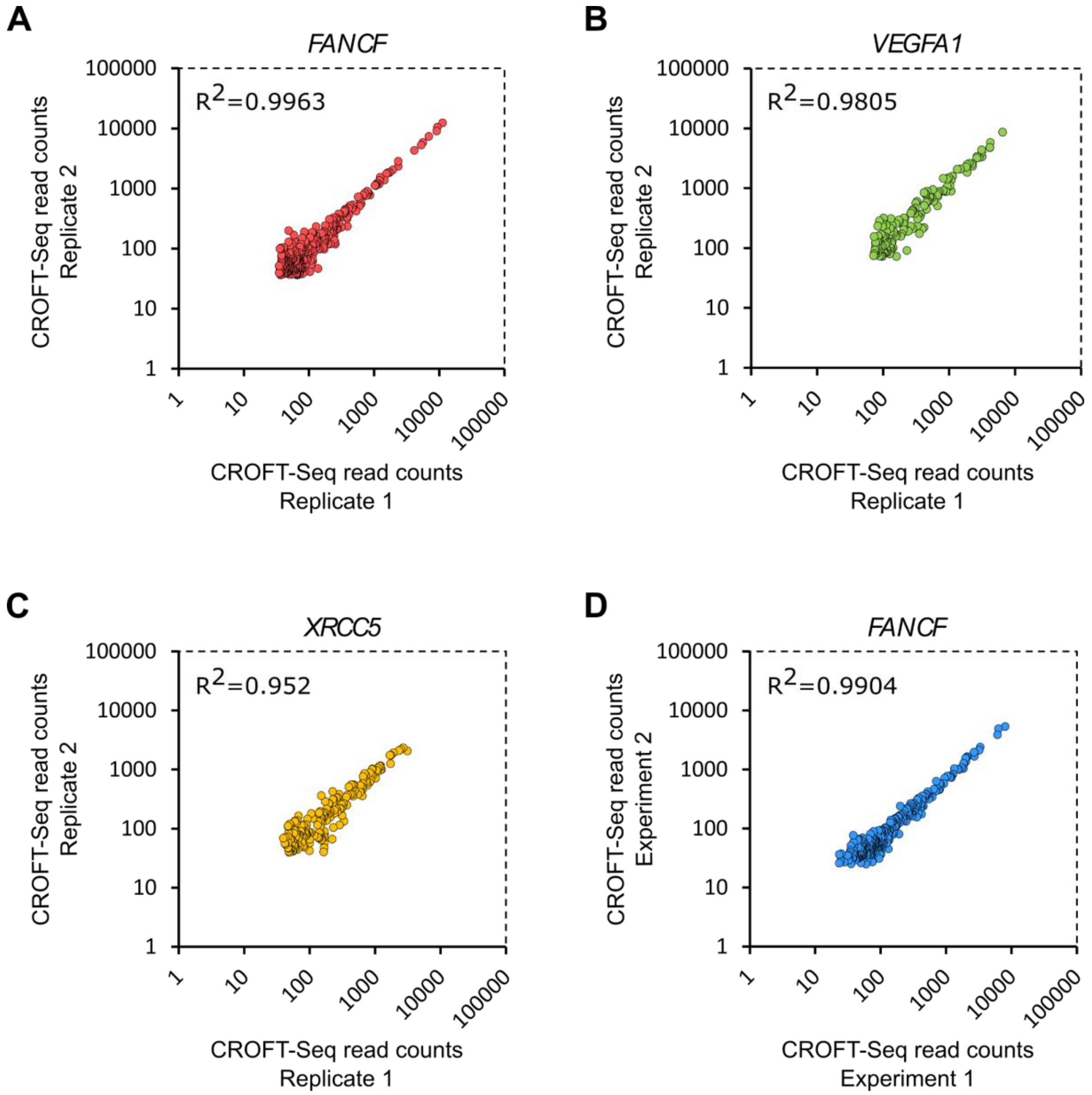


**Supplementary Figure S2.** Reproducibility of CROFT-Seq read counts. (**A-C**) Scatterplots showing a read count (log scale) correlation between two independent CROFT-Seq technical replicates, with gRNA targeting the *FANCF*, *VEGFA1,* and *XRCC5* sites. For data visualization, the top1000 sites of each replicate were analyzed and plotted only those found in both replicates. (**D**) Scatterplot showing read count correlation between two independent CROFT-Seq biochemical experiments (each reaction is an average of three technical replicates) where gRNA is targeted against the *FANCF* site. For data visualization, the top1000 sites of each replicate were analyzed and plotted only those found in both replicates.

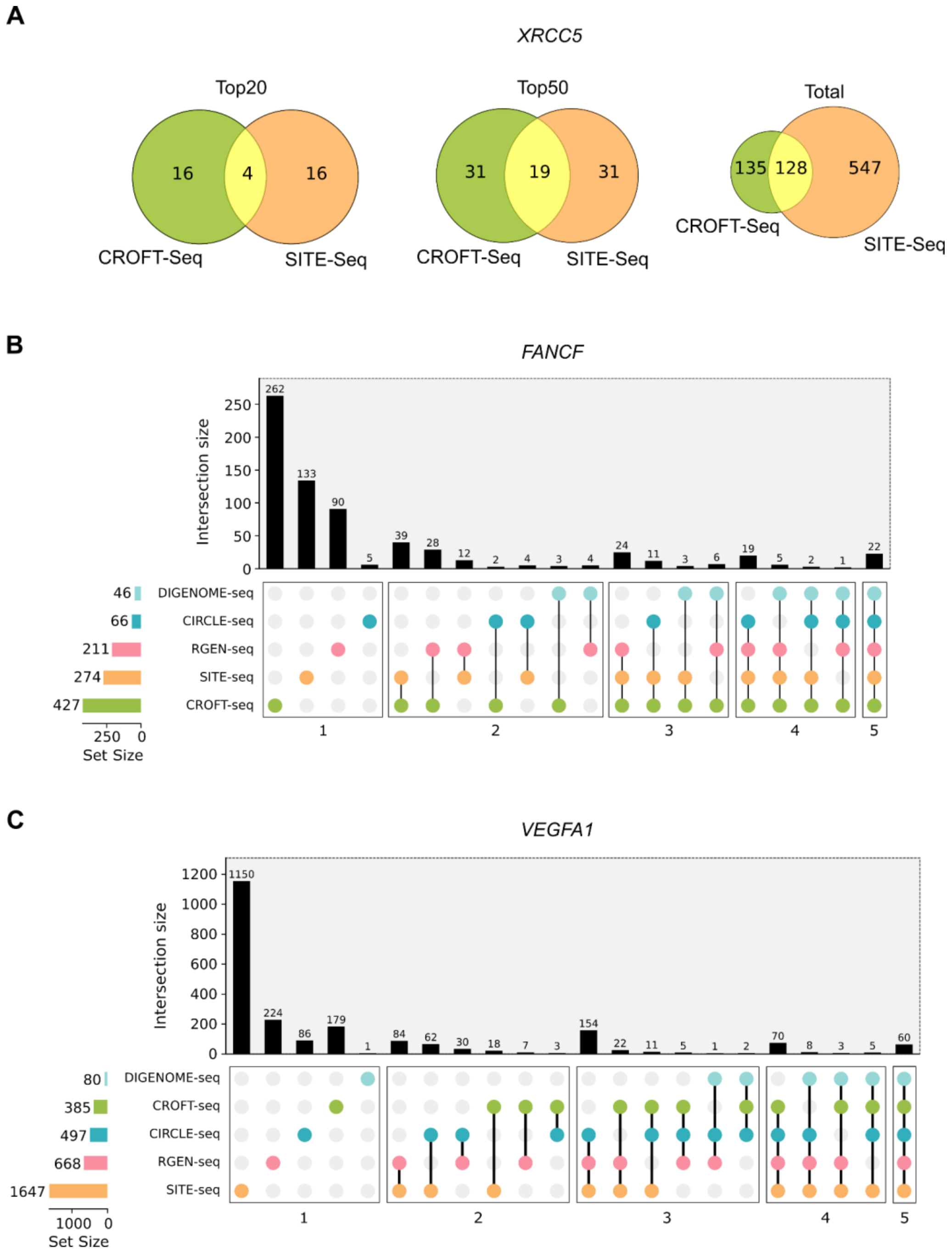


**Supplementary Figure S3.** Comparison of CROFT-Seq with other *in vitro* methods for off-target detection. (**A**) Venn diagrams showing overlap at different ranges (top20, top50 and whole off-target list) of detected off-target sites between CROFT-Seq and SITE-Seq using gRNA targeting the *XRCC5* site. (**B**) Upset plot showing the overlap of detected off-target sites between CROFT-Seq, DIGENOME-Seq, CIRCLE-Seq, RGEN-Seq, and SITE-Seq using gRNA targeting the *FANCF* site. The total number of off-target sites detected is shown on the left. The number of off-target methods detecting the same off-target sites is shown at the bottom. (**C**) Upset plot showing the overlap of detected off-target sites between CROFT-Seq, DIGENOME-Seq, CIRCLE-Seq, RGEN-Seq, and SITE-Seq using gRNA targeting the *VEGFA1* site. The total number of off-target sites detected is shown on the left. The number of off-target methods detecting the same off-target sites is shown at the bottom.

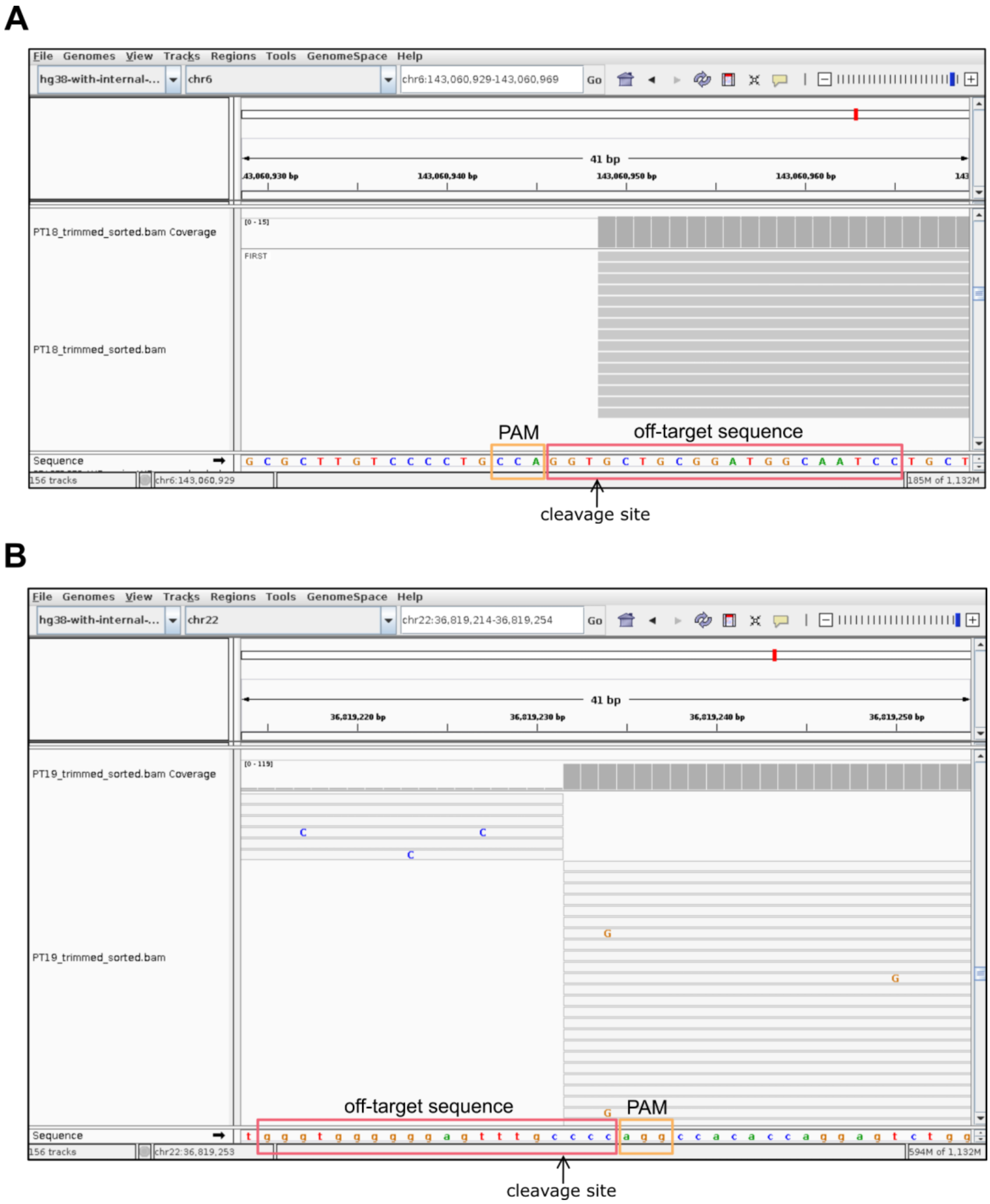


**Supplementary Figure S4.** GUIDE-Seq off-target sites not detected by the CROFT-Seq. IGV snapshots of the *FANCF* (**A**) and *VEGFA1* (**B**) off-target sites, which were not detected by the CROFT-Seq. Read alignment views at a base-pair resolution are shown in a 41 bp window. The coverage plots represent average summarized reads. Aligned reads represent the cleavage site of the *FANCF* off-target site (5'-GGATTGCCATCCGCAGCACCTGG-3') and the *VEGFA1* off-target site (5'-GGGTGGGGGGAGTTTGCCCCAGG-3') not detected by the CROFT-Seq.
